# BayCount: A Bayesian Decomposition Method for Inferring Tumor Heterogeneity using RNA-Seq Counts

**DOI:** 10.1101/218511

**Authors:** Fangzheng Xie, Mingyuan Zhou, Yanxun Xu

## Abstract

Tumors are heterogeneous - a tumor sample usually consists of a set of subclones with distinct transcriptional profiles and potentially different degrees of aggressiveness and responses to drugs. Understanding tumor heterogeneity is therefore critical for precise cancer prognosis and treatment. In this paper, we introduce BayCount, a Bayesian decomposition method to infer tumor heterogeneity with highly over-dispersed RNA sequencing count data. Using negative binomial factor analysis, BayCount takes into account both the between-sample and gene-specific random effects on raw counts of sequencing reads mapped to each gene. For the posterior inference, we develop an efficient compound Poisson based blocked Gibbs sampler. Simulation studies show that BayCount is able to accurately estimate the subclonal inference, including number of subclones, the proportions of these subclones in each tumor sample, and the gene expression profiles in each subclone. For real-world data examples, we apply BayCount to The Cancer Genome Atlas lung cancer and kidney cancer RNA sequencing count data and obtain biologically interpretable results. Our method represents the first effort in characterizing tumor heterogeneity using RNA sequencing count data that simultaneously removes the need of normalizing the counts, achieves statistical robustness, and obtains biologically/clinically meaningful insights. The R package BayCount implementing our model and algorithm is available for download.

## 1 Introduction

Tumor heterogeneity (TH) is a phenomenon that describes distinct molecular profiles of different cells in one or more tumor samples. TH arises during the formation of a tumor as a fraction of cells acquire and accumulate different somatic events (*e.g.*, mutations in different cancer genes), resulting in heterogeneity within the same biological tissue sample and between different ones, spatially and temporally (Russnes et al., 2011; Ding et al., 2012). As a result, tumor cell populations are composed of different subclones (subpopulations) of cells, characterized by distinct genomes, transcriptional profiles (Kim et al., 2015), as well as other molecular profiles, such as copy number alterations. Understanding TH is critical for precise cancer prognosis and treatment. Heterogenetic tumors may exhibit different degrees of aggressiveness and responses to drugs among different samples due to genetic or gene expression differences. The level of heterogeneity itself can be used as a biomarker to predict treatment response or prognosis since more heterogeneous tumors are more likely to contain treatment-resistant subclones (Marusyk et al., 2012). This will ultimately facilitate the rational design of combination treatments, with each distinct compound targeting a specific tumor subclone based on its transcriptional profile.

Large-scale sequencing techniques provide valuable information for understanding tumor complexity and open a door for the desired statistical inference on TH. Previous studies have focused on reconstructing the subclonal composition by quantifying the structural subclonal copy number variations (Carter et al., 2012; Oesper et al., 2013), somatic mutations (Nik-Zainal et al., 2012; Roth et al., 2014; Xu et al., 2015), or both (Deshwar et al., 2015; Lee et al., 2016). In this paper, we aim to learn tumor transcriptional heterogeneity using RNA sequencing (RNA-Seq) data.

In the analysis of gene expression data, matrix decomposition models have been extensively studied in the context of microarray and *normalized* RNA-Seq data (Venet et al., 2001; Lähdesmäki et al., 2005; Wang et al., 2006; Abbas et al., 2009; Repsilber et al., 2010; Shen-Orr et al., 2010; Gong et al., 2011; Hore et al., 2016; Wang et al., 2016). Generally, given gene expression data matrix *X* = (*x_ij_*)_*G×S*_, where the (*i, j*)th element records the expression value of the *i*th gene in the *j*th sample, they decompose *X* by modeling *x_ij_* with 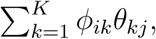 where *ϕ_ik_* encodes the expression level of the *i*th gene in the *k*th subclone, *_kj_* represents the mixing weight of the *k*th subclone in the *j*th sample, and *K* is the number of subclones. The decomposition can be solved by either optimization algorithms (Venet et al., 2001; Wang et al., 2016) or statistical inference by assuming a normal distribution on *x_ij_*. While it is reasonable to assume normality for microarray gene expression data, it is often inappropriate to adopt such an assumption for directly modeling RNA-Seq data, which involve nonnegative integer observations. If a model based on normal distribution is used, one often needs to first normalize RNA-Seq data before performing any downstream analysis. See Dillies et al. (2013) for a review on normalization methods. Although normalization often destroys the nonnegative and discrete nature of the RNA-Seq data, it remains the predominant way for data preprocessing due to not only the computational convenience in modeling normalized data, but also the lack of appropriate count data models. Distinct from previously proposed methods, in this paper, we propose an attractive class of count data models in decomposing RNA-Seq count matrices.

There are, nevertheless, statistical challenges with RNA-Seq count data. First, the distributions of the RNA-Seq count data are typically over-dispersed and sparse. Second, the scales of the read counts in sequencing data across samples can be enormously different due to the mechanism of the sequencing experiment such as the variations in technical lane capacities. The larger the library sizes (*i.e.*, sequencing depths) are, the larger the read counts tend to be. In addition, the differences in gene lengths or GC-content (Pickrell et al., 2010) can bias gene differential expression analysis, particularly for lowly expressed genes (Oshlack and Wakefield, 2009). A number of count data models have been developed for RNA-Seq data (Lee et al., 2013; Kharchenko et al., 2014; Fan et al., 2016). For example, Lee et al. (2013) proposed a Poisson factor model on microRNA to reduce the dimension of count data and identify low-dimensional features, followed by a clustering procedure over tumor samples. Kharchenko et al. (2014) developed a method using a mixture of negative binomial and Poisson distributions to model single cell RNA-Seq data for gene differential expression analysis. None of these methods, however, address the problem of TH.

To this end, we propose BayCount, a Bayesian matrix decomposition model built upon the negative binomial model (Zhou, 2016), to infer tumor transcriptional heterogeneity using RNA-Seq count data. BayCount accounts for both the between-sample and genespecific random effects and infers the number of latent subclones, the proportions of these subclones in each sample, and subclone-specific gene expression simultaneously. The R package BayCount implementing our model and algorithm is available at http://pages.jh.edu/~fxie5/Research/BayCount_0.1.0.tar.gz with the installation script at http://pages.jh.edu/~fxie5/Research/Installation_script.R.

The remainder of the paper is organized as follows. In Section 2, we introduce BayCount, a hierarchical Bayesian model for RNA-Seq count data, and develop an e cient compound Poisson based blocked Gibbs sampler. We investigate the performance of the posterior inference and robustness of BayCount through extensive simulation studies in Section 3, and apply BayCount to analyze two real-world RNA-Seq datasets from The Cancer Genome Atlas (TCGA) (Cancer Genome Atlas Research Network, 2012) in Section 4. We conclude the paper in Section 5.

## 2 Hierarchical Bayesian Model and Inference

In this section we present the proposed hierarchical model for RNA-Seq count data, develop the corresponding posterior inference, and discuss how to determine the number of subclones.

### 2.1 BayCount Model

We assume that *S* tumor samples are available from the same or different patients. Consider a *G × S* count matrix *Y* = (*y_ij_*)_*G×S*_, where each row represents a gene, each column represents a tumor sample, and the element *y_ij_* records the read count of the *i*th gene from the *j*th tumor sample. The Poisson distribution Pois(λ) with mean *λ* > 0 is commonly used for modeling count data. Poisson factor analysis (PFA) (Zhou et al., 2012) factorizes the count matrix *Y* as *y_ij_* ∼ Pois 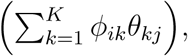, where 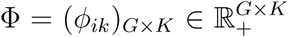 is the factor loading matrix and 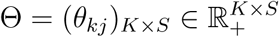 is the factor score matrix. Here *K* is an integer indicating the number of latent factors, and each column of Φ is subject to the constraint that 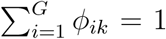 and *Φ_ij_* ≥ 0. However, the restrictive equidispersion property of the Poisson distribution that the variance and mean are the same limits the application of PFA in modeling sequencing data, which are often highly over-dispersed. For this reason, one may consider negative binomial factor analysis (NBFA) of Zhou (2016) that factorizes *Y* as *y_ij_* ∼ NB 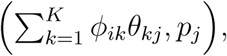 where *p_j_* ∈ (0, 1). We denote *y* ∼ NB(*r, p*) as a negative binomial distribution with shape parameter *r* > 0 and success probability *p* ∈ (0, 1), whose mean and variance are *rp*/(1 − *p*) and *rp*/(1 − *p*)^2^, respectively, with the variance-to-mean ratio as 1/(1 − *p*).

Denote the *j*th column of *Y* as *y_j_* = (*y*_1*j*_, *y*_2_*j*, &, *y_Gj_*)^*T*^, the count profile of the *j*th tumor sample. To account for both the between-sample and gene-specific random effects when modeling RNA-Seq count data, we propose

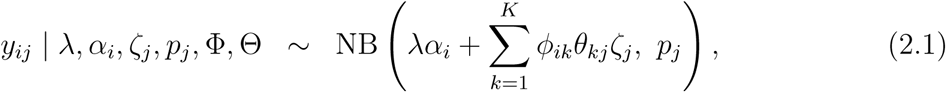

where *_i_* accounts for the gene-specific random effect of the *i*th gene, and *p_j_* control the overall scale of the gene-specific effects and between-sample effect of the *j*th sample, respectively, and 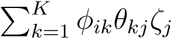 represents the average effect of the *K* subclones on the expression of the *i*th gene in the *j*th sample.

To see this, recall that the mean of *y_ij_* based on (2.1) is

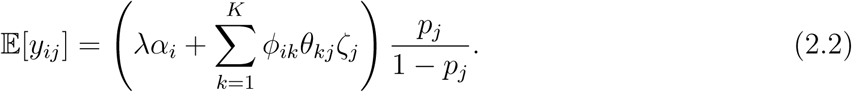

Since *p_j_* is sample-specific, the term *p_j_*/(1 − *p_j_*) describes the effect of sample *j* on read counts due to technical or biological reasons (*e.g.*, different library sizes, biopsy sites, etc). We assume the relative expression of the *i*th gene in the *k*th subclone is described by *ϕ_ik_*, where *ϕ_ik_* ≥ 0. Since the sample-specific effect has already been captured by *p_j_*, for modeling convenience, we normalize the gene expression so that the expression levels sum to one for each subclone. Namely, 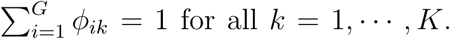 Furthermore, we assume that *θ_kj_* represents the proportion of the *k*th subclone in the *j*th sample, where *θ_kj_* ≥ 0 and 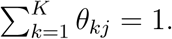 We can interpret *θ_kj_ζ_j_* as the population frequency of the *k*th subclone in the *j*th sample, where parameter *ζ_j_* controls the scale. Together, the summation 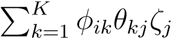 represents the aggregated expression level of the *i*th gene across all *K* subclones in the *j*th sample. To further account for the gene-specific random effects that are independent of the samples and subclones, we introduce an additional term *_i_* to describe the random effect of the *i*th gene on the read counts such as GC-content and gene length. We assume 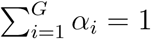 so that *α_i_* represents the relative gene-specific random effect of the *i*th gene with respect to all the genes and controls the overall scale of the gene-specific random effects.

Following Zhou (2016), the model in (2.1) has an augmented representation as

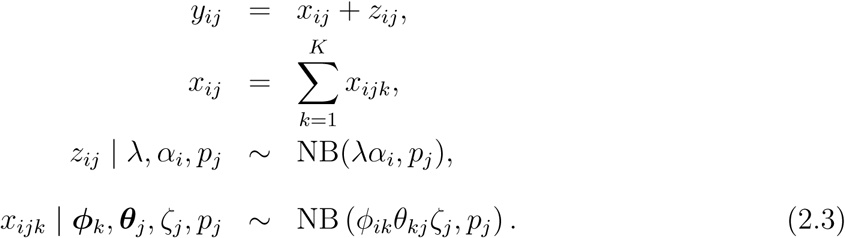

From (2.3), the raw count *y_ij_* of the *i*th gene in the *j*th sample can be interpreted as coming from multiple sources: *x_ijk_* represents the count of the *i*th gene contributed by the *k*th subclone in the *j*th sample, where *k* = 1*, &, K*, while *z_ij_* is the count contributed by the gene-specific random effect of the *i*th gene in the *j*th sample. Note that for the auxiliary count matrix (*x_ij_*)_*G×S*_, we factorize the negative binomial shape parameter matrix into the product of and under the negative binomial likelihood. This is different from the exponential family formulation of non-negative matrix factorization in Ghahramani et al. (2014), as the negative binomial distribution NB(*r, p*) belongs to the exponential family only if the shape parameter *r* is fixed.

Denote 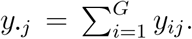 Since 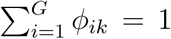 and 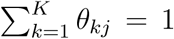 by construction, under (2.3), by the additive property of independent negative binomial random variables with the same success probability, we have

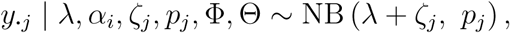

and, in particular, the mean as E[*y_·j_*] = (*λ + ζ_j_*)*p_j_*/(1 − *p_j_*) and the variance as Var(*y_·j_*) = E[*y_·j_*] + E^2^[*y_·j_*]/(λ + *ζ_j_*). It is clear that *p_j_*, the between-sample random effect of the *j*th sample, governs the variance-to-mean ratio of *y_·j_*, whereas λ + *ζ_j_*, the sum of the scale λ of the gene-specific random effects and the scale *ζ_j_* for the *j*th sample, controls the quadratic relationship between Var(*y_·j_*) and E[*y_·j_*].

We complete the model by setting the following priors that will be shown to be amenable to the posterior inference:

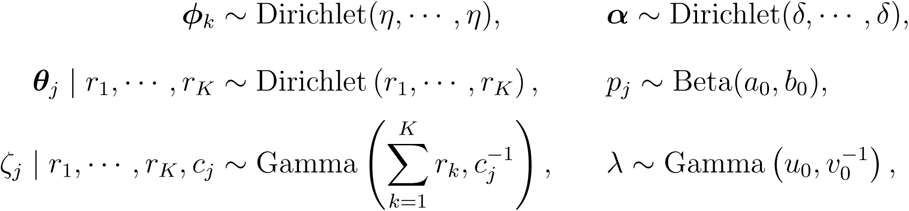

where *ϕ_k_* = (*ϕ*_1*k*_, …, *ϕ_Gk_*)^*T*^, *θ_j_* = (*θ*_1*j*_, …, *θ_Kj_*)^*T*^, *α* = (*α*_1_, …, *α_G_*)^*T*^, Gamma(*a, b*) denotes a gamma distribution with mean *ab* and variance *ab*^2^, and Dirichlet(*η*_1_, …, *η_d_*) denotes a *d*-dimensional Dirichlet distribution with parameter vector (*η*_1_, …, *η_d_*). We further impose the hyperpriors, expressed as *r_k_* | γ_0_, *c*_0_ ∼ Gamma 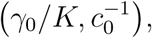 *c_j_* ∼ Gamma 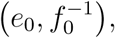 γ_0_ ∼ Gamma 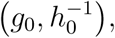 and *c_0_* ∼ Gamma 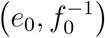 to construct a more flexible model. Shown in Figure 1 is the graphical representation of BayCount.

**Figure 1:**
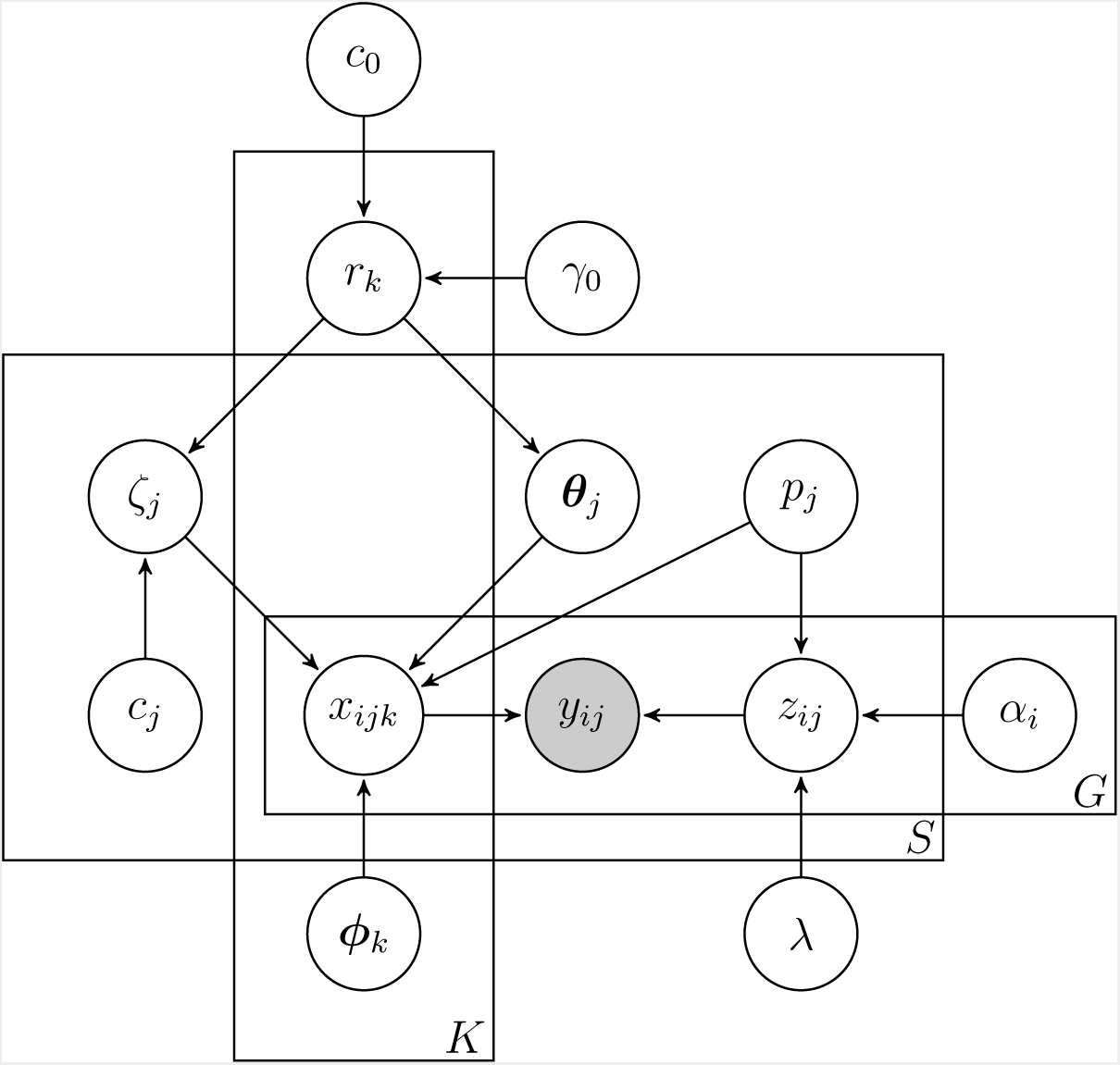
Graphical representation of BayCount. The boxes represent replicates. For example, the box containing *r_k_, x_ijk_* and *ϕ_k_*, with *K* in its bottom right corner, indicates that there are *K* “copies” of *r_k_, x_ijk_* and *ϕ_k_* with *k* = 1, …, *K*. Shaded nodes represent observations.

### 2.2 Gibbs Sampling via Data Augmentation

For BayCount, while the full conditional posterior distributions of *p_j_*, *c_j_* and *c*_o_ are straightforward to derive due to conjugacy, a variety of data augmentation techniques are used to derive the closed-form Gibbs sampling update equations for all the other model parameters. Rather than going into the details here, let us first assume that we have already sampled the latent counts *x_ijk_* given the observations *y_ij_* and model parameters, which, according to Theorem 1 of Zhou (2016), can be realized by sampling from the Dirichlet-multinomial distribution. Given *x_ijk_*, we derive the Gibbs sampling update equations for Φ and Θ via data augmentation. Then we describe in Section A of the Supplementary Material a compound Poisson based blocked Gibbs sampler that completely removes the need of sampling *x_ijk_*.

**Sampling Φ and Θ**

We introduce an auxiliary variable *ℓ_ijk_* that follows a Chinese restaurant table (CRT) distribution, denoted by *ℓ_ijk_* | *x_ijk_, ϕ_ik_θ_kj_ζ_j_* ∼ CRT(*x_ijk_, ϕ_ik_θ_kj_ζ_j_*), with probability mass function

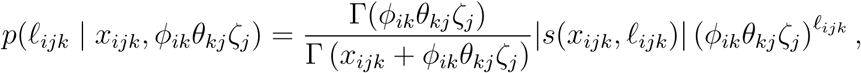

supported on {0, 1, 2, …, *x_ijk_*}, where *s*(*x_ijk_, ℓ_ijk_*) are Stirling numbers of the first kind (Johnson et al., 1997). Sampling *ℓ* ∼ CRT(*x, r*) can be realized by taking the summation of *m* independent Bernoulli random variables: 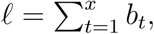 where *b_t_* ∼ Bernoulli (*r*/(*r* + *t* − 1)) independently. Following Zhou and Carin (2012), the joint distribution of *ℓ_ij_* and *x_ij_* described by

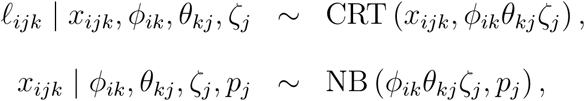

can be equivalently characterized under the compound Poisson representation

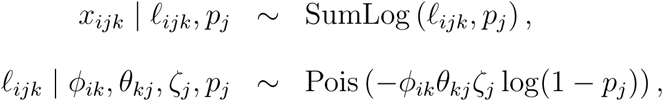

where *x* ∼ SumLog (*ℓ, p*) denotes the sum-logarithmic distribution generated as 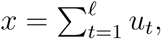 where 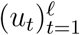 are independent, and identically distributed (i.i.d.) according to the logarithmic distribution (Quenouille, 1949) with probability mass function *p*(*u*) = *−p^u^*/[*u* log(1 − *p*)], supported on {1, 2, …}.

Under this augmentation, the likelihood of *ϕ_ik_*, *θ_kj_* and *ζ_j_* becomes

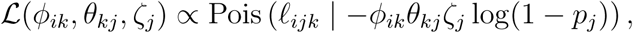

where Pois(· | *λ*) denotes the probability mass function of the Poisson distribution with mean *λ*. It follows immediately that the full conditional posterior distributions for *ϕ_k_* and *θ_j_* are

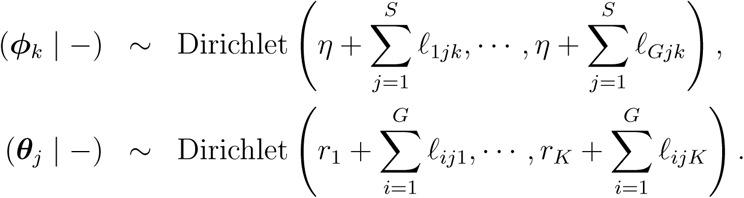

Using data augmentation, we can similarly derive the full conditional posterior distributions for *ζ_j_*, *α*, *r_k_* and *γ*_0_, as described in detail in Section A of the Supplementary Material.

### 2.3 Determining the Number of Subclones *K*

We have so far assumed *a priori* that *K* is fixed. Determining the number of factors in factor analysis is, in general, challenging. Zhou (2016) suggested adaptively truncating *K* during Gibbs sampling iterations. This adaptive truncation procedure, which is designed to fit the data well, may tend to choose a large number of factors, some of which may be highly correlated to each other and hence are potentially redundant. To facilitate the interpretation of the model output, we seek a model selection procedure that estimates *K* in a more conservative manner. It is critical to select a moderate *K* that is large enough to fit the data reasonably well, but at the same time is small enough for the sake of interpretation. As is suggested by Shen and Huang (2008) and Ghahramani et al. (2014), one way of determining *K* is cross-validation. This method is computationally expensive since it requires repeated leave-out testing procedure for each fixed *K*.

Alternatively, one can generalize the idea of “finding the elbow of scree plots”, an ad-hoc method for selecting the latent dimension in principal component analysis (Zhu and Ghodsi, 2006). In the scree plots, the reductions of the residual sum of squares, a measurement of goodness-of-fit to the data, are plotted against the latent dimension. An “elbow” is the point that maximizes the difference of the slopes of the two adjacent line segments. Generalizing to BayCount, we calculate the estimated log-likelihood of the model under different numbers of subclones (using post-burn-in MCMC samples) as the measurement of the goodness-of-fit to the data. These samples are obtained by running the compound Poisson based blocked Gibbs sampler for different *K*’s. The estimate of *K* is the point at which an apparent decrease in the slopes of segments that connect the log-likelihood log *L*(*K*) evaluated at two consecutive *K* values is detected. Formally, we denote the log-likelihood function log *L*(*K*) as a function of *K*, and define the second-order finite difference Δ^2^log *L*(*K*) of the log-likelihood function by Δ^2^log *L*(*K*) := 2 log *L*(*K*) − log *L*(*K* − 1) − log *L*(*K* + 1), for *K* = *K*_min_ + 1, …, *K*_max_ − 1, where *K*_min_ and *K*_max_ are the lower and upper limits of *K*, respectively. Then an estimate of *K* is given by *K̂* = arg max_*K*_ Δ^2^log *L*(*K*). Notice that similar approaches are adopted to detect the number of latent factors in the context of time series of inhomogeneous Poisson processes (Shen and Huang, 2008) and Poisson factor models (Lee et al., 2013).

## 3 Simulation Study

In this section, we evaluate BayCount through simulation studies. Two different scenarios are considered.

- Scenario I. We simulate the data according to BayCount itself in (2.1). In particular, we generate the subclone-specific gene expression data matrix = (*ϕ_ik_*)_*G×K*_ ∈ 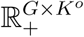 by i.i.d. draws of *ϕ_k_* ∼ Dirichlet(0.05, …, 0.05), the proportion matrix Θ = 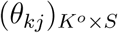 by i.i.d. draws of *_j_* ∼ Dirichlet(0.5, …, 0.5), and *ζ_j_* by i.i.d. draws of *ζ_j_* ∼ Gamma(0.5*K°,* 1), where *i* = 1, …, *G*, *j* = 1, …, *S*, and *k* = 1, …, *K°*. Here *G* is the number of genes, *S* is the number of samples, and *K°* is the simulated number of subclones. We set = 1, draw from Dirichlet(0.5, …, 0.5), and generate *p_j_* from a uniform distribution such that the variance-to-mean ratio *p_j_*/(1 − *p_j_*) of *y_·j_* ranges from 100 to 10^6^, encouraging the simulated data to be over-dispersed.
- Scenario II. To evaluate the robustness of BayCount, we simulate the data from a model that is different from BayCount. We generate the subclone-specific gene expression data matrix 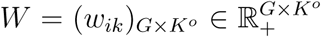 by i.i.d. draws of *W_ik_* ∼ Gamma(0.05, 10), and the proportion matrix Θ = (*θ_kj_*)_*K*°×S_ by i.i.d. draws of *θ_j_* ∼ Dirichlet(0.5, …, 0.5). We set λ = 1, draw from Dirichlet(0.5, …, 0.5), and generate *p_j_* from a uniform distribution such that the variance-to-mean ratio *p_j_*/(1 − *p_j_*) of *y*_·_j ranges from 100 to 10^6^. The count matrix *Y* = (*y_ij_*)_*G×S*_ is generated from 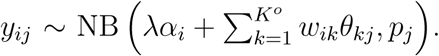 Note that in scenario II the scales of *W* = (*w_ik_*)_*G×K*°_ are not subject to the constraint

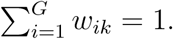

We will show that BayCount can accurately recover both the subclone-specific gene expression patterns and subclonal proportions. The hyperparameters are set to be *η* = 0.1, *a*_0_ = *b*_0_ = 0.01, *e*_0_ = *f*_0_ = 1, *g*_0_ = *h*_0_ = 1, and *u*_0_ = *v*_0_ = 100. We consider *K* ∈ {2, 3, …, 10}. The compound Poisson based blocked Gibbs sampler is implemented with an initial burn-in of *B* = 1000 iterations, followed by *n* = 1000 post-burn-in iterations. Notice that for any permutation matrix Π ∈ {0, 1}*^K×K^*, ΦΠ^T^ΠΘ = ΠΘ, leading to potential label switching phenomenon during the MCMC. The following procedure is implemented in practice to address this issue.

- **Step 1:** Collect the *n* post-burn-in MCMC samples 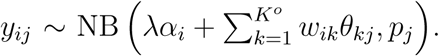
- **Step 2:** Find the posterior MCMC sample that maximizes the log-likelihood:
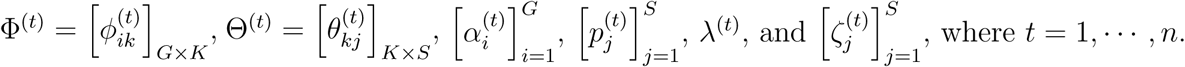
- **Step 3:** For *t* = 1, 2, …, n, find 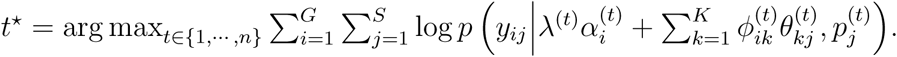 where the arg min is taken over all *K × K* permutation matrices and ‖ · ‖*_F_* is the matrix Frobenius norm.
- **Step 4:** For *t* = 1, 2, …, *n*, replace Φ^(*t*)^ by Φ^(*t*)^Π^(*t*)T^ and Θ^(*t*)^ by Π^(*t*)^Θ^(*t*)^.

After implementing the procedure above for the posterior samples of and, we compute the posterior means and 95% credible intervals for all parameters using the post-burn-in MCMC samples.

### 3.1 Synthetic data with *K°* = 3

We first simulate two datasets with *G* = 100, *S* = 20, and *K°* = 3 under both scenario I and scenario II. Under scenario I, the data generation scheme is the same as BayCount. Figure S1 in the Supplementary Material plots Δ^2^log *L*(*K*) versus *K*, indicating *K̂* = 3, which is the same as the simulation truth. In terms of estimating *K*, we also compare BayCount with three alternative competitors: Bayesian information criterion (BIC), deviance information criterion (DIC), and the logarithmic conditional predictive ordinate (log-CPO). See Section C of the Supplementary Material for the detailed results and comparisons of estimating *K*. The estimated subclone-specific gene expression matrix Φ̂ and subclonal proportions Θ̂ are computed as the posterior means of the post-burn-in MCMC samples. Figure S2 and S3 compare the simulated true Φ and Θ with the estimate Φ̂ and Θ̂, respectively. We can see that both the subclone-specific gene expression patterns and the subclonal proportions are successfully recovered.

A competitive alternative for RNA-seq decomposition is the non-negative matrix factorization (NMF) (Lee and Seung, 1999) to the normalized expression data. The normalized expression data are obtained by taking the Anscombe transformation (Anscombe, 1948) to the original count matrix: 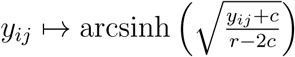 for some constants *r* and *c*, *i* = 1, …, *G*, *j* = 1, …, *S*. The detailed results and comparison between BayCount and the NMF on the normalized expression data are provided in Section E of the Supplementary Material. As shown in Figures S20-22, BayCount outperforms the NMF in terms of estimating the number of subclones, the subclonal expression, and the subclonal proportions.

The analysis under scenario II is of greater interest, since the focus is to evaluate the robustness of BayCount. BayCount yields an estimate of *K̂* = 3, as shown in Figure S4. We then focus on the posterior inference based on *K̂* = 3. Figure 2 compares the estimated subclonal proportions Θ̂ with the simulated true subclonal proportions across samples, along with the posterior 95% credible intervals. The results show that the estimate Θ̂ approximates the simulated true Θ well. We then report the posterior inference on the subclone-specific gene expression Φ. Notice that under BayCount, 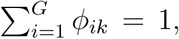 and hence the estimate Φ̂ by BayCount and the unnormalized gene expression profile matrix *W* used in generating the simulated data are not directly comparable. To see whether the gene expression patterns are recovered, we first normalize *W* by its column sums as *Ŵ* = *W*Λ^−1^, where Λ = diag 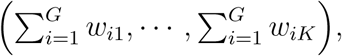, so that *ŵ_ik_* represents the relative expression level of the *i*th gene in the *k*th subclone, and then compare ̂̂ with Ŵ. For visualization, the genes with small standard deviations (less than 0.01) are filtered out due to their indistinguishable expressions across different subclones. Figure 3 compares the heatmap of Φ̂, with the heatmap of the simulated true (normalized) subclone-specific gene expression *Ŵ* on selected differentially expressed genes. It is clear that the patterns of subclone-specific gene expression estimated under BayCount closely match the simulation truth. We also evaluate the stability of BayCount by adding independent Pois(10) noisy counts to the original count matrix as perturbations and then analyze the perturbed count matrix using BayCount. Figures S18 and S19 in Section D.2 of the Supplementary Material indicate that BayCount is stable in the presence of noisy perturbations.

**Figure 2:**
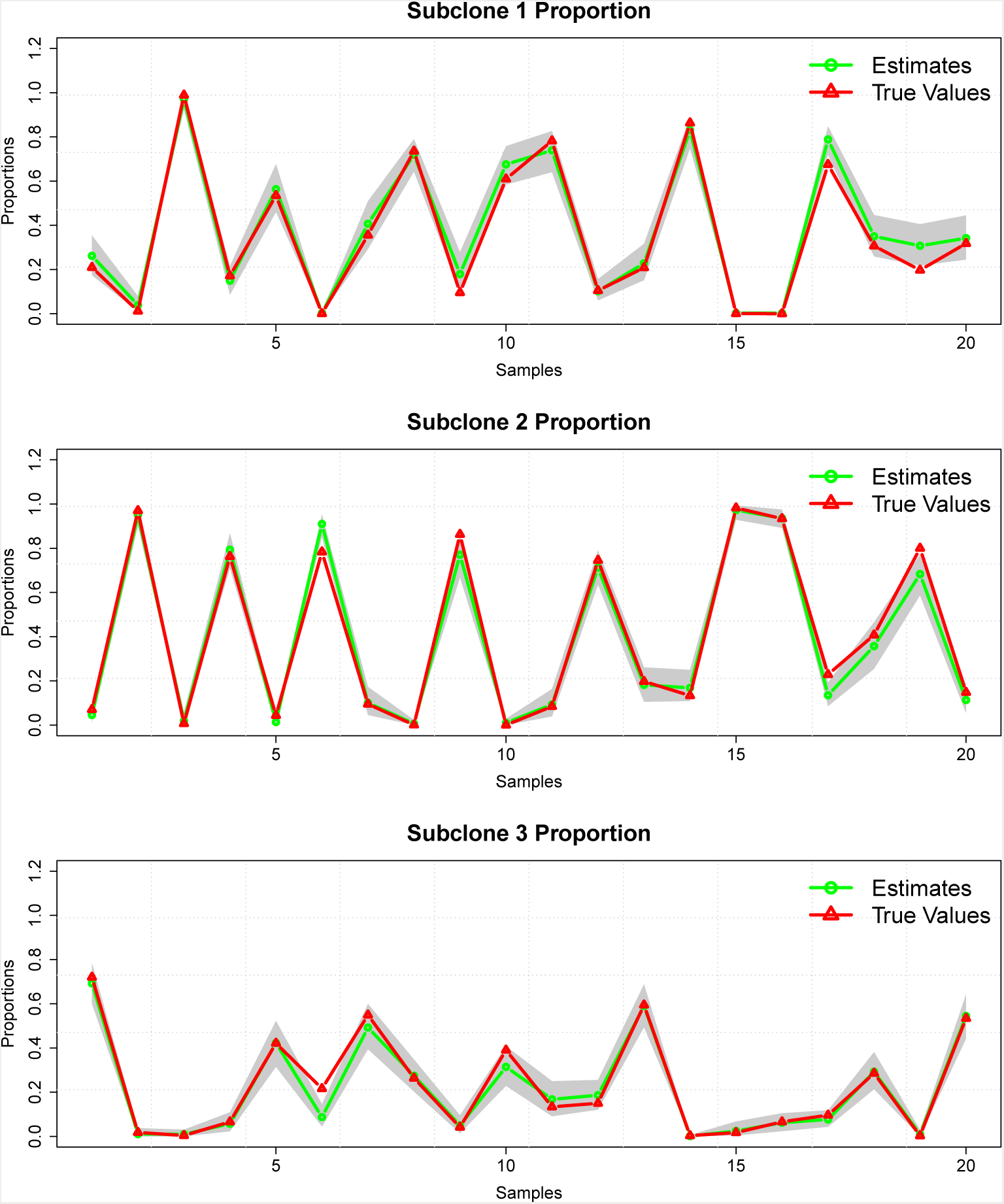
Figure 2: The estimated subclonal proportions Θ̂ across samples *j* = 1, …, 20 for the synthetic dataset with *K°* = 3 under scenario II. Horizontal axis is the index *j* = 1, …, 20 of tumor samples, and vertical axis is the proportion. The green lines represent the estimate Θ̂, and the red lines represent the simulated true subclonal proportions. The shaded area represents the posterior 95% credible bands.

**Figure 3:**
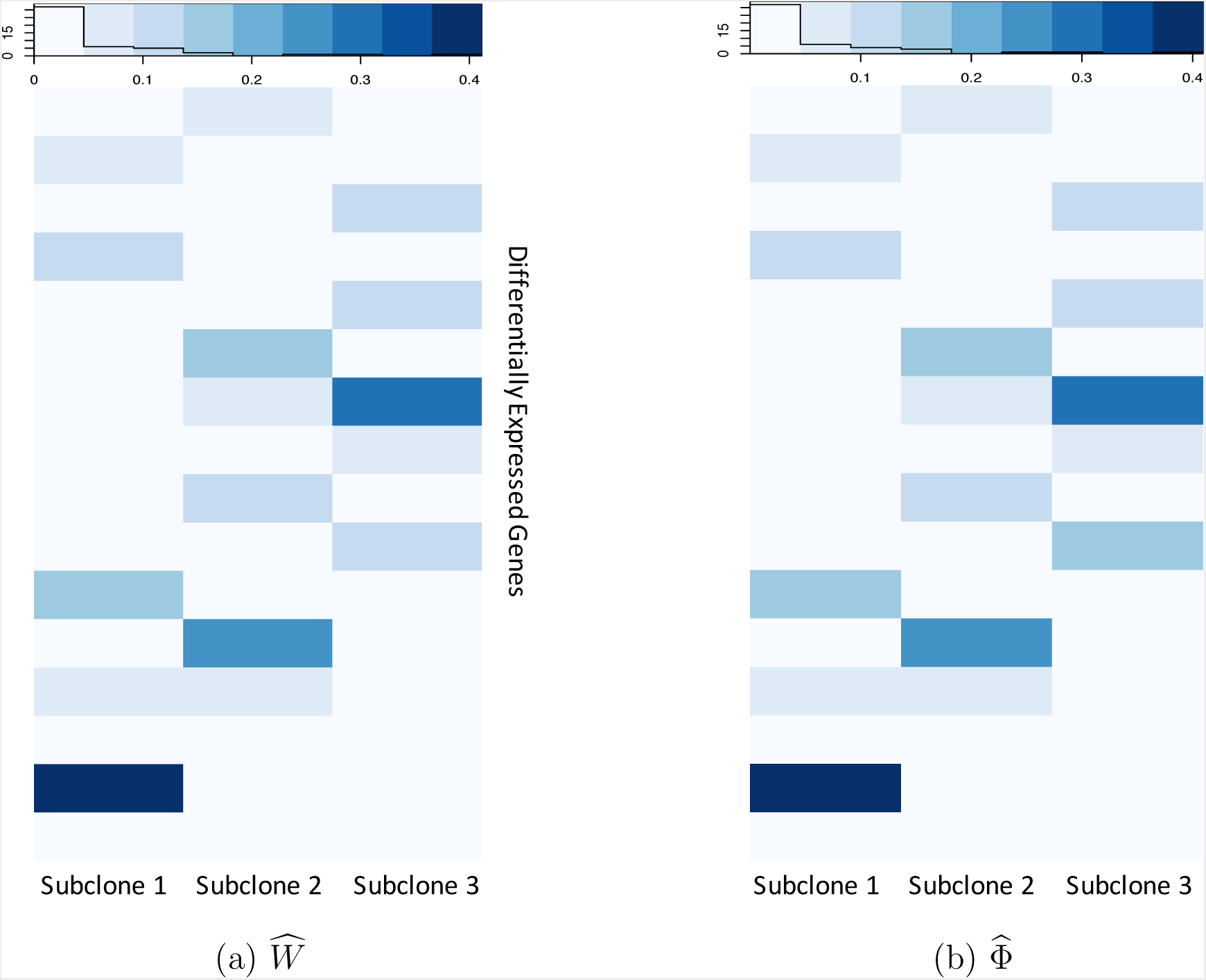
Comparison of subclone-specific gene expression patterns for the synthetic dataset with *K°* = 3 under scenario II. Panel (a) is the heatmap of *Ŵ*, computed by normalizing the simulated true expression data *W* by its column sums, and panel (b) is the heatmap of the estimate Φ̂.

### 3.2 Synthetic data with *K°* = 5

Similarly as in Section 3.1, we simulate two datasets with *G* = 1000, *S* = 40, and *K°* = 5 under scenarios I and II, respectively. Under scenario I, BayCount yields an estimate of *K̂* = 5 (Figure S5), and from Figures S6 and S7, both the subclone-specific gene expression patterns and the subclonal proportions are successfully captured.

Under scenario II, BayCount yields an estimate of *K̂* = 5 (Figure S8). For the subclonal proportions = (*_kj_*)_*K×S*_, Figure 4 shows that the estimate Θ̂ successfully recovers the simulated true proportions. Notice that the credible bands are narrower than those in Figure 2, implying relatively smaller uncertainty in estimating subclonal proportions for larger dataset. Figure S9 presents the autocorrelation plots of the posterior samples of some randomly selected *_kj_*’s generated by the compound Poisson based blocked Gibbs sampler, indicating that the Markov chains mix well. The Markov chains also mix well when the sample size *S* varies over {40, 80, 120, 200}, where the synthetic dataset is simulated with *G* = 1000 and *K* = 5 under scenario II. See Section F Figure S25 in the Supplementary Material for the trace plots of some randomly selected *_kj_*’s when *S* varies.

**Figure 4:**
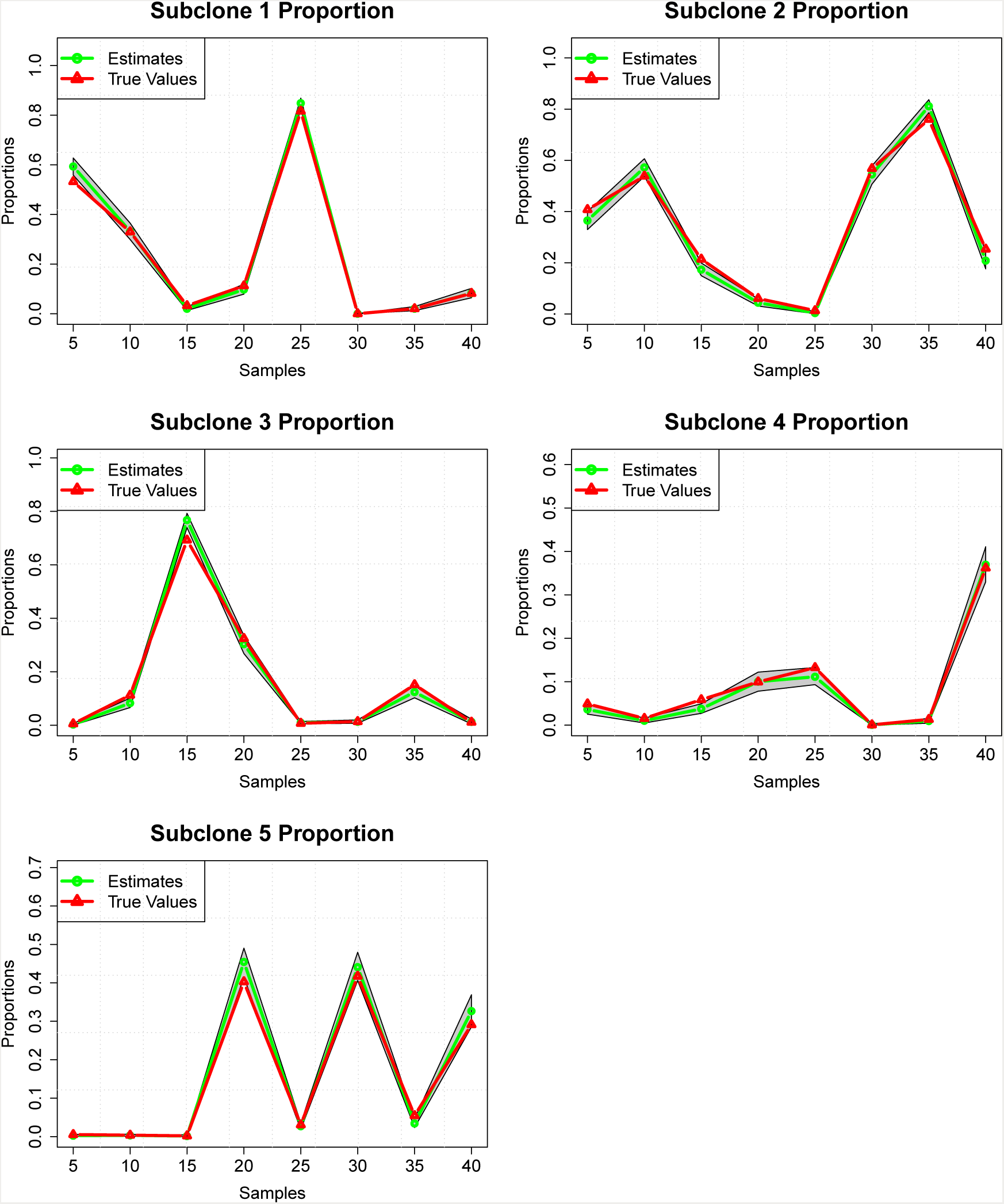
Subclonal proportions across samples *j* = 5, 10, …, 35, 40 for the synthetic dataset with *K°* = 5 under scenario II. Horizontal axis is the index of tumor samples, and vertical axis is the proportion. The green lines represent Θ̂, and red lines represent the simulated true subclonal proportions. The shaded area represents the posterior 95% credible bands.

Figure 5 compares the simulated true (normalized) subclone-specific gene expression *Ŵ* with the estimate Θ̂ under BayCount. For this dataset we pre-screen *Ŵ* with a threshold 0.008 on the across-subclone standard deviations for all genes for visualization. The high concordance between the heatmaps of the estimated and true expression patterns of the differentially expressed genes indicates that the subclone-specific gene expression patterns have been successfully recovered as well.

**Figure 5:**
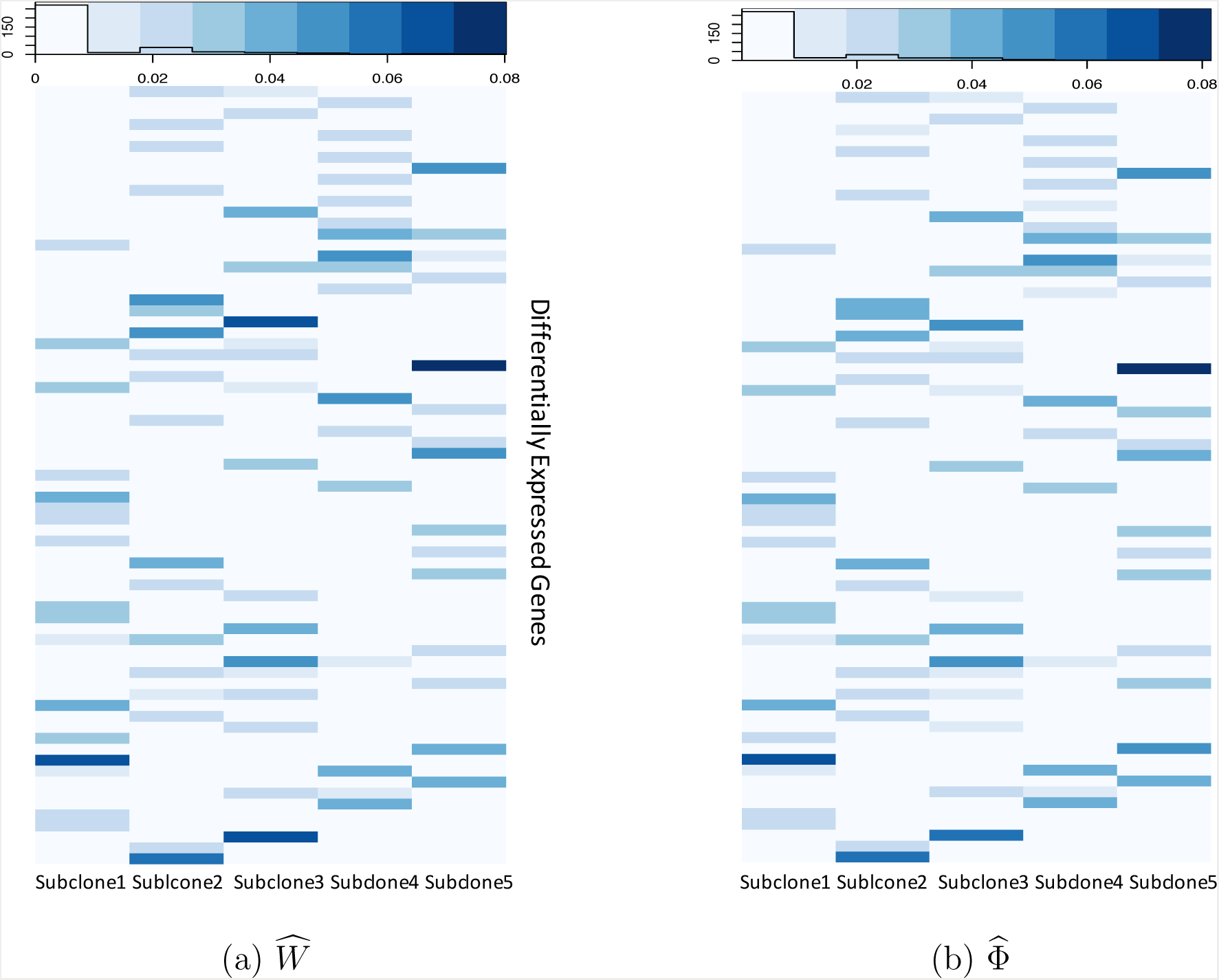
Comparison of subclone-specific gene expression patterns for the synthetic dataset with *K°* = 5 under scenario II. Panel (a) is the heatmap of *Ŵ*, computed by normalizing the simulated true expression data *W* by its column sums, and panel (b) is the heatmap of the estimate Φ̂.

In summary, BayCount can accurately identify the number of subclones, estimate the subclonal proportions in each sample, and recover the subclone-specific gene expression patterns of the differentially expressed genes.

## 4 Real-world Data Analysis

We implement and evaluate BayCount on the RNA-Seq data from The Cancer Genome Atlas (TCGA) (Cancer Genome Atlas Research Network, 2012) to study tumor heterogeneity (TH) in both lung squamous cell carcinoma (LUSC) and kidney renal clear cell carcinoma (KIRC). We first run the proposed Gibbs sampler for each fixed *K* ∈ {2, 3, …, 10}, compute both the posterior mean log *L*(*K*) of the log-likelihood for each fixed *K*, and estimate *K* by maximizing Δ^2^log *L*(*K*) over *K*. Next, based on the estimate *K̂* and the posterior samples generated by the proposed Gibbs sampler, we estimate the proportions of the identified subclones in each tumor sample and the subclone-specific gene expression, which in turn can be used for a variety of downstream analyses.

### 4.1 TCGA LUSC Data Analysis

We apply BayCount to the TCGA RNA-Seq data in lung squamous cell carcinoma (LUSC), which is a common type of lung cancer that causes nearly one million deaths worldwide every year. We downloaded FASTQ formatted files for LUSC tumor samples via the National Cancer Institute’s Cancer Genomics Hub (Wilks et al., 2014) and then used the featureCounts function in the Rsubread package (Liao et al., 2014; Rahman et al., 2015) to obtain integer-based, gene-level read counts. We select 200 primary tumor samples and 382 previously reported important lung cancer genes (Wilkerson et al., 2010; Cancer Genome Atlas Research Network, 2012) for analysis of LUSC, such as KRAS, STK11, BRAF, and RIT1.

BayCount yields an estimate of five subclones (Figure S10) and their proportions in each tumor sample are shown in Figure 6. To identify the dominant subclone for each sample, we compare the estimate Θ̂ of the five subclones in each tumor sample, and use them to cluster the patients. Formally, for each patient *j* = 1, …, *S*, we compute the dominant subclone 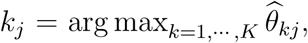 and then cluster patients according to {*j* : *k_j_* = *k*}, *k* = 1, &, *K̂*. That is to say, the patients with the same dominant subclone belong to the same cluster. We next check if the identified subclones have any clinical utility, *e.g.*, stratification of patients in terms of overall survival. Figure 7a shows the Kaplan-Meier plots of the overall survival of the patients among the five clusters identified by their dominant subclones. Indeed, patients stratified by these five BayCount-identified groups exhibit very distinct survival patterns (log-rank test *p* value = 0.0194).

**Figure 6:**
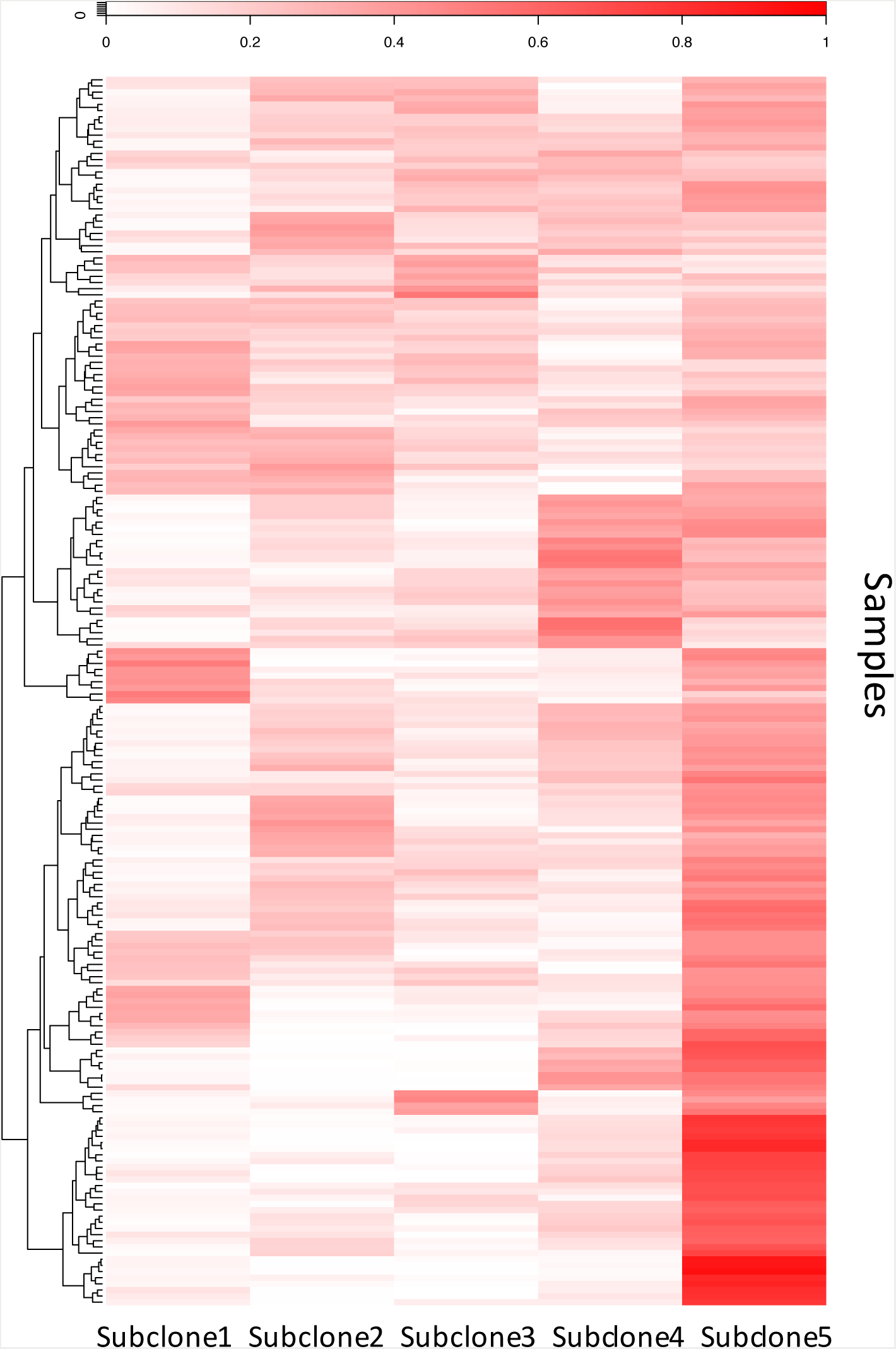
Heatmap of the subclonal proportions across LUSC tumor samples *j* = 1, …, 200. From the heatmap it is clear that subclone 5 occupies relatively larger proportions for a large number of patients than the other 4 subclones.

**Figure 7:**
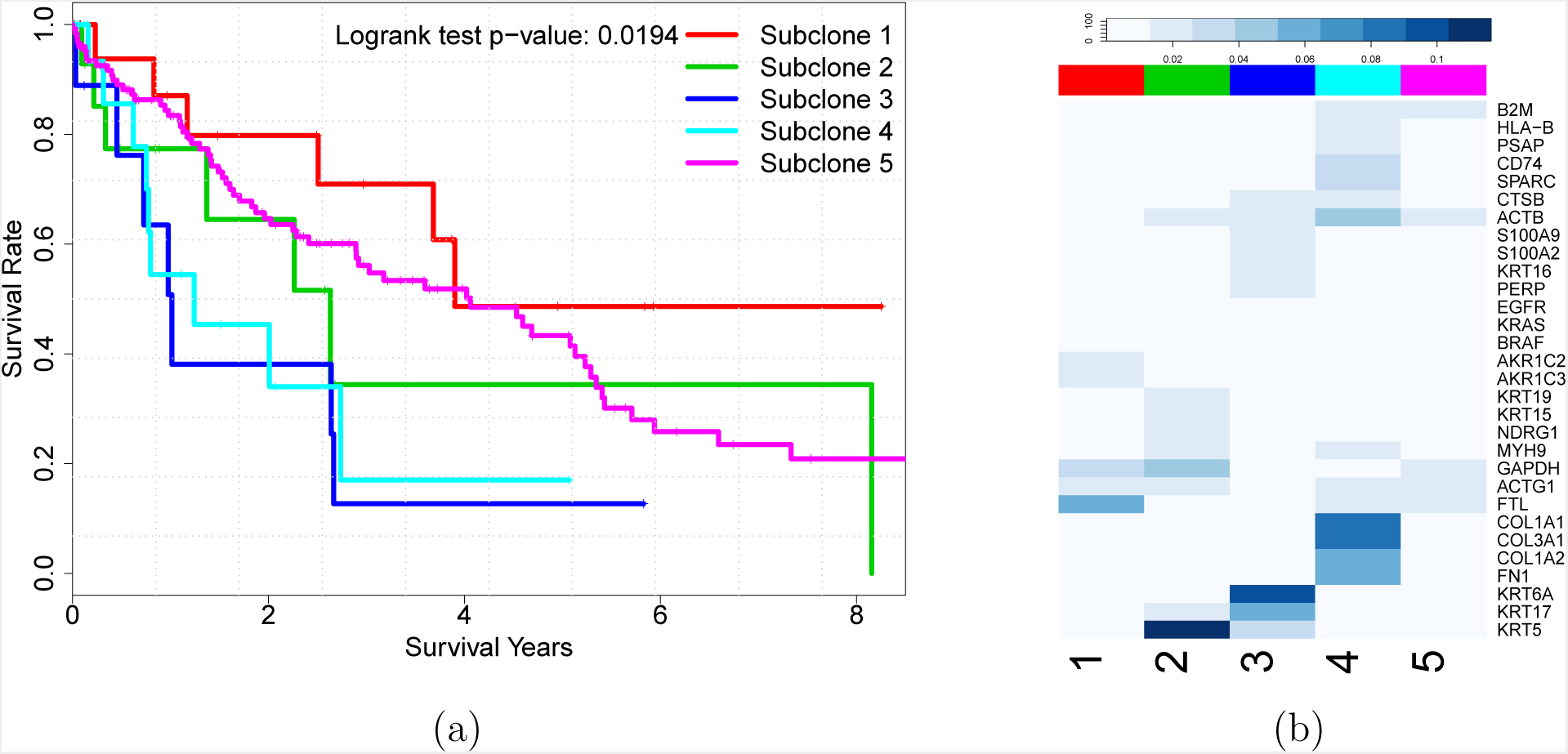
Panel (a) shows the Kaplan-Meier plots of overall survival in the LUSC dataset, where the patients are stratified by five clusters identified by subclone domination under BayCount. Panel (b) shows the subclone-specific gene expression of the top differentially expressed genes among five subclones.

For comparison, we implement the NMF to the normalized expression data after Anscombe by the two NMF-identified groups do not exhibit distinct survival patterns (log-rank test *p* value = 0.125, see Figure S24a). The detailed results and comparisons are provided in Section E of the Supplementary Material.

Figure 7b shows the expression levels of the top 30 differentially expressed genes (ranked by the standard deviations of the subclone-specific gene expression levels *ϕ_ik_*’s in an increasing order) in these five subclones. Distinct expression patterns are observed among different subclones. For example, the FTL level is elevated in subclone 1; the expression levels of several genes encoding keratins (KRT5, KRT6A, etc.) are elevated in subclone 3; and the COL1A1 and COL1A2 expression levels are elevated in subclone 4. Interestingly, the patients with these dominant subclones also show the expected survival patterns. The subclone-1 dominated patients have better overall survival. Previous studies show that the expression of FTL is decreased in lung tumors compared to normal tissues (Kudriavtseva et al., 2009), and one plausible explanation is that subclone 1 may descend from less malignant cells and therefore resemble (or consist of) normal cells. Keratins and collagen I (encoded by COL1A1 and COL1A2) are known to play key roles in epithelial-to-mesenchymal transition (EMT), which subsequently initiates metastasis and promotes tumor progression (DePianto et al., 2010; Karantza, 2011; Shintani et al., 2008). This agrees with our observation of worse prognosis in patients who have either subclone 3 (with elevated Keratin-coding genes) or subclone 4 (with elevated collagen I coding genes) as their dominant subclone.

With the inferred 5 subclones and the corresponding parameters of the LUSC dataset under BayCount, we perform an additional simulation study that is realistic: we simulate a synthetic dataset using the parameters inferred from the LUSC dataset with *G* = 382, *S* = 200, and ground true *K* = 5. Figures S15-17 show that BayCount successfully recovers the ground true *K*, the subclonal expression patterns, and the subclonal proportions. See Section D.1 of the Supplementary Material for the detailed results.

### 4.2 Kidney Cancer (KIRC) Data Analysis

Similarly, we obtain gene level read counts (Liao et al., 2014) for 200 TCGA kidney renal clear cell carcinoma (KIRC) tumor RNA-seq samples and analyze them under BayCount. Among a total of 23,368 genes, 966 significantly mutated genes (Cancer Genome Atlas Research Network, 2013) in KIRC patients are selected, including VHL, PTEN, MTOR, etc.

BayCount yields an estimate of five subclones in KIRC (Figure S11). Figure 8 shows the Kaplan-Meier plots of the overall survival of the patients grouped by their dominant subclones (panel a) and the heatmap of the subclone-specific gene expression corresponding to the top 30 differentially expressed genes (panel b). Since we have a large number of genes to begin with, whereas 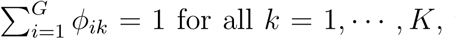 the subclone-specific gene expression estimates Φ̂ will be small. For better visualization, we plot Φ̂ in the logarithmic scale. The subclonal proportions across 200 KIRC tumor samples are shown in Figure S12. As shown in Figure 8, the patients with these dominant subclones again show distinct survival patterns. One of the poor survival groups (dominated by subclone 5) is characterized by elevated expression of TGFBI, which is known to be associated with poor prognosis (Zhu et al., 2015) and matches our observation here.

**Figure 8:**
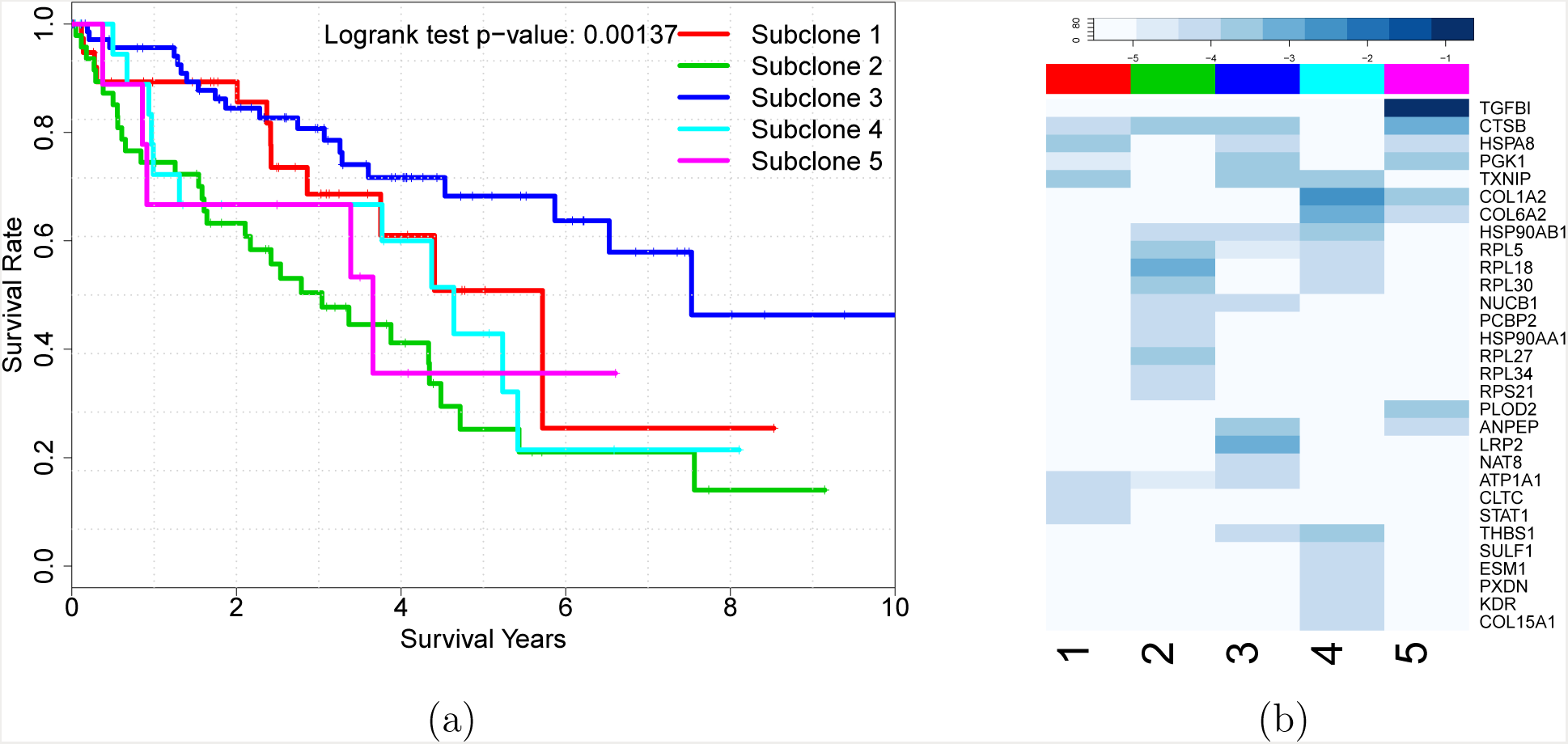
Panel (a) shows the Kaplan-Meier plots of overall survival in the KIRC dataset, where the patients are stratified by five clusters identified by subclone domination under BayCount. Panel (b) shows the subclone-specific gene expression (in the logarithmic scale) of the top differentially expressed genes among the five inferred subclones.

One distinction of our method from conventional subgroup analysis methods is that we focus on characterizing the underlying subclones (*i.e.*, biologically meaningful subpopulations), by not only their individual molecular profiles but also their proportions. Instead of grouping the patients by their dominant subclones, we examine the proportion itself in terms of clinical utility. Interestingly, as shown in Figure 9a, the proportion of subclone 2 increases with tumor stage: *i.e.*, as subclone 2 expands and eventually outgrows other subclones, the tumor becomes more aggressive. In contrast, the proportion of subclone 3 decreases with tumor stage (Figure 9b). Subclone 3 might be characterized by the less malignant (or normal-like) cells and takes more proportion in the beginning of the tumor life cycle. As tumor progresses to more advanced stages, subclone 3 could be suppressed by more aggressive subclones (*e.g.*, subclone 2) and takes a decreasing proportion. Unsurprisingly, the survival patterns agree with our speculations about subclones 2 and 3, with the patients dominated by subclone 2 (the more aggressive subclone) and subcolone 3 (the less aggressive subclone) showing the worst and best survivals, respectively.

**Figure 9:**
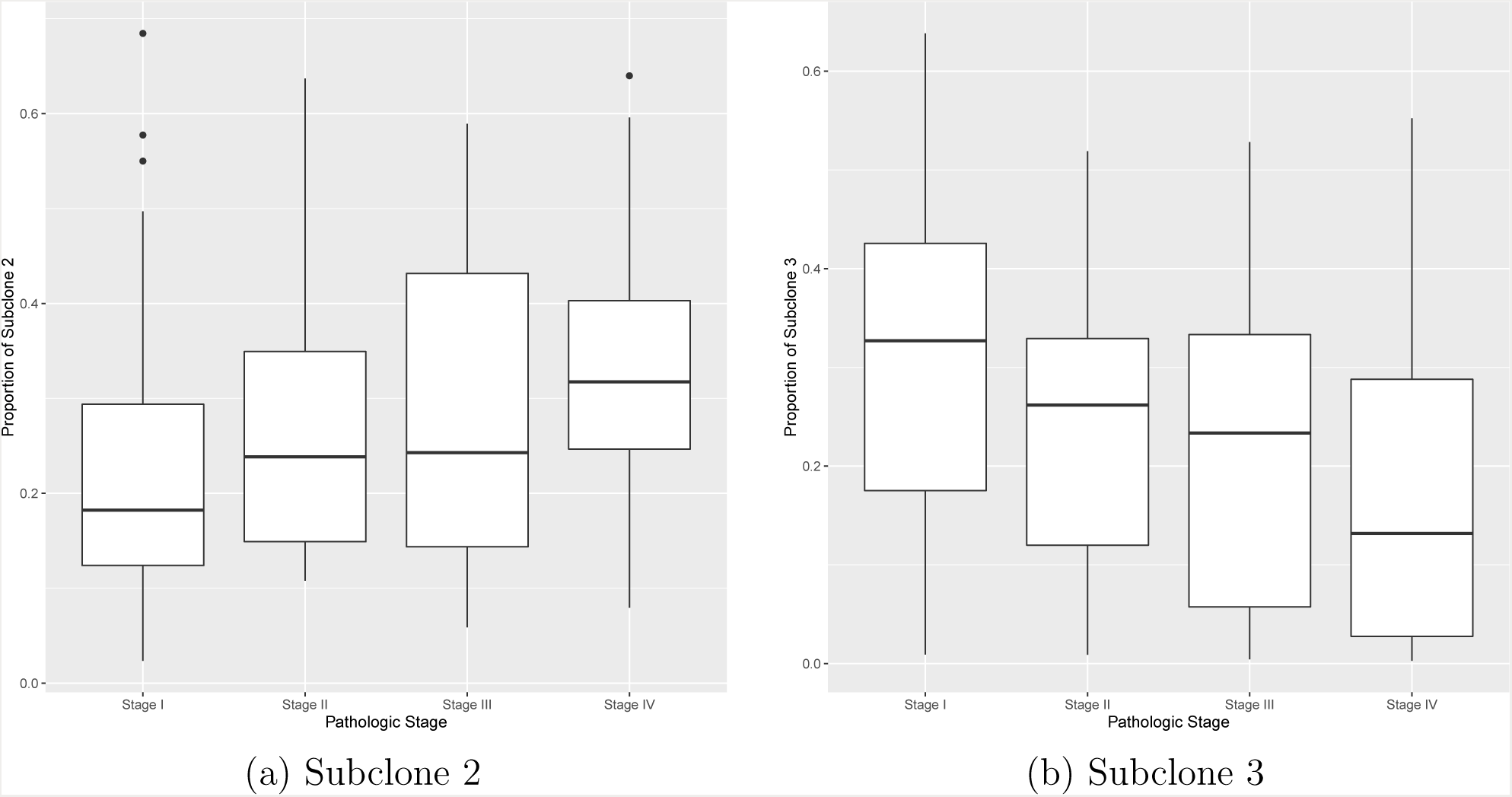
Panel (a): the proportions of subclone 2 in each tumor sample versus their pathologic stages (*p*-value = 0.00173). Panel (b): the proportions of subclone 3 in each tumor sample versus their pathologic stages (*p*-value = 0.00299).

More excitingly, we find that the proportions of these two subclones can complement clinical variables in further stratifying patients. For patients at early stage where the event rate is low and clinical information is relatively limited, the proportions of subclones 2 and 3 serve as a potent factor in further stratifying patients (Figure S13) when dichotomizing at a natural cutoff . Combining our observations above, subclone proportions may provide additional insights into the progression course of tumors, assistance in biological interpretation, and potentially more accurate clinical prognosis.

## 5 Conclusion

The emerging high-throughput sequencing technology provides us with massive information for understanding tumors’ complex microenvironment and allows us to develop novel statistical models for inferring tumor heterogeneity. Instead of normalizing RNA-Seq data that may bias downstream analysis, we propose BayCount to directly analyze the raw RNA-Seq count data. Overcoming the natural challenges of analyzing raw RNA-seq count data, BayCount is able to factorize them while adjusting for both the between-sample and gene-specific random effects. Simulation studies show that BayCount can accurately recover the subclonal inference used to generate the simulated data. We apply BayCount to the TCGA LUSC and KIRC datasets, followed by correlating the subclonal inferences with their clinical utilities for comparison. In particular, by grouping patients according to their dominant subclones, we observe distinct and biologically sensible overall survival patterns for both LUSC and KIRC patients. Moreover, the proportions of the subclones may complement clinical variables in further stratifying patients. In addition to prognosis value, tumor heterogeneity may be used as a biomarker to predict treatment response. For example, tumor samples with large proportions of cells bearing higher expressions on clinically actionable genes should be treated differently from those that have no or a small proportion of such cells. In addition, metastatic or recurrent tumors may possess very different compositions of subclones and should be treated differently.

BayCount provides a general framework for inference on latent structures arising naturally in many other biomedical applications involving count data. For example, analyzing single-cell data is a potential further application of BayCount due to their sparsity and over-dispersion nature. Macosko et al. (2015) describe Drop-Seq, a technology for profiling more than 40,000 single cells at one time. The unique characteristic of dropped-out events (Fan et al., 2016) in single cell sequencing limits the applicability of normalization methods in bulk RNA-Seq data. Also, such huge amount number of single-cells and high levels of sparsity pose di culties for dimensionality reduction methods such as principal component analysis. Inferring distinct cell populations in single-cell RNA count data will be an interesting extension of BayCount.

## Acknowledgement

Yanxun Xu’s research is partly supported by Johns Hopkins *in* Health and Booz Allen Hamilton.

